# Effect of sleeve gastrectomy on sperm quality and fertility on non-obese diabetes diet model rat

**DOI:** 10.1101/2023.11.08.566186

**Authors:** Gamze Tumentemur, Mustafa Titiz, Alev Bobus Ors

## Abstract

**Background:** In people with diabetes, the effect of sleeve gastrectomy on impaired sperm parameters, hormonal profile and testis tissue remains controversial to some extent.

**The context and purpose of the study:** Effects of sleeve gastrectomy on hormonal profile, sperm parameters, and testis tissue in infertile men with type II diabetes mellitus (TIIDM). This study included thirty two TIIDM that were scheduled with and without sleeve gastrectomy. All rats were taken a sperm analysis, a testis tissue and a serum hormone profile before, 8 weeks after surgery.

**Results:** There was a significant correlation between weight loss after sleeve gastrectomy and decreased in glucose profile (p < 0.05). As regards the hormonal profile, testosterone improved significantly after 8 weeks following sleeve gastrectomy. There was a significant increase in sperm count (p < 0.05), and improved sperm morphology during the follow-up after sleeve gastrectomy, also showed that significant changes as improved in testis tissue after surgery.

**Conclusion:** Sleeve gastrectomy significantly improves testosterone deficiency, testis tissue and sperm count in TIIDM rats. More prospective clinical studies are needed to show how bariatric surgery affects infertility affected by TIIDM patients.

## Introduction

Metabolic Syndrome is the most important component as known TIIDM, is become a problem for all countries in the world [1]. In recent years, TIIDM has also created complications on male reproductive system functions [2, 3, 4, 5]. Recent reports have shown that different molecular mechanisms, such as altered reproductive hormone levels, neuropathy, and increased oxidative stress, in diabetic men, are responsible for the structural damage and dysfunction of sperm by affecting the continuity of spermatogenesis [2]. By affecting the hypothalamic-pituitary gonadal axis of TIIDM, it affects the glucose mechanisms of testicular and sperm cells, reducing the levels of luteinizing hormone (LH) and follicle stimulating hormone (FSH) identified in the plasma of diabetic patients [5]. The management and treatment of complications continue to be a priority for governments around the world because only the economic burden it brought in 2007 exceeded $ 174 billion [6]. There are also clinical trials conducted in parallel with experimental studies [7] showing testosterone, LH, and FSH levels in diabetes [6]. A decrease in testosterone level is one of the causes of diabetic testicular damage. Many researchers suggest that the decrease in serum testosterone level in TIIDM is due to steroidogenetic damage in Leydig cells [8]. It is known that insulin resistance and increased insulin levels suppress LH, SHBG, and testosterone level, while bariatric surgery also reduces insulin resistance and insulin levels [9,10]. Bariatric surgery is currently emerging as an area suitable for establishing surgical procedures not only for obesity but especially for diabetes treatment [11]. It provides more successful results in the treatment of other diseases accompanying TIIDM than medical treatment [12, 13] There are many methods developed for obesity and surgical treatment of TIIDM in the last 30 years. While some of these methods have been abandoned over time, some of them continue to be applied successfully thanks to their effective long-term results. According to the American Metabolic and Bariatric Surgery Association, 179,000 bariatric surgery operations carried out in America in 2013, and there has been a 15% increase in bariatric surgery operations since 2011 [14]. Chair of the Health Technology Assessment (2014) “The Role of Obesity Treatment of Obesity Surgery in Turkey” has performed bariatric surgery in the number of cases reported in Turkey; It has been given as 2197 in 2008, 3268 in 2010 and 4511 in 2012, and the number of new obesity cases is increasing gradually. There are three bariatric surgery methods most commonly used in Turkey; sleeve gastrectomy, mini gastric bypass, and gastric band [15]. Sleeve gastrectomy, also called tube stomach, is a bariatric surgery technique that has been increasing in use in recent years and effective in a short time (16, 17). The most popular surgical procedure for treating morbid obesity is laparoscopic sleeve gastrectomy; however, if dietary changes are not made afterward, the fertility damage caused by a high-fat diet and TIIDM will persist. The effects of male erectile function are controversial due to the contradiction in sperm parameters along with testosterone level after bariatric operations. In this study, a rat non-obese model of TIIDM was developed and the effects on male fertility with and without sleeve gastrectomy were evaluated.

## Research Design and Methods

### Animals and Ethics Statement

3-month-old male Sprague–Dawley rats (n=32) weighing 300-350 gr were purchased and treated in accordance with the Guide for the Care and Use of Laboratory Animals (ACUDEHAM 2018-06). All rats housed individually in ventilated cages at a constant 24–26 °C temperature and humidity, and a 12-h light/dark cycle. Rats with BMI results over 5 kg / m^2^ were considered obese and were not included in the study. Non obese rats were induced a single dose of 60 mg/kg streptozotocin (STZ) (Sigma-Aldrich, catalog number; S0130) dissolved in 10 mmol/l citrate buffer (pH 4.5) was intraperitoneally injected to rats. Random glucose was consecutively measured by monitoring proximal tail vein for blood glucose measurement using a glucometer (Accu-Chek II Boehringer Mannheim Canada, Dorval, Quebec). Rats with more than three random glucose level measurements > 250 mg/dl were considered diabetic. Sprague Dawley male rats were divided into four groups; while rats in group 1 remained on the regular rat chow (no procedure was applied for 8 weeks), group 2 received a dose of STZ to induce diabetes (no procedure was applied for 8 weeks) (18,19), group 3 received a dose of STZ to induce diabetes and created sham), group 4 received a dose of STZ to induce diabetes and created sleeve gastrectomy (the rats were anesthetized and the lower left part of the diaphragm was incised and the stomach was made visible. Starting immediately distal to the stomach incisura angularis, most of the curvature major of the stomach was removed, starting with a linear stapler and the stomach was retracted laterally. Then, the sleeve gastrectomy method was applied with linear staples sequential firings towards the angle of His. Animals were fasted for 24 hours prior to surgery, and (Ketalar 90 mg/k (ketamine) +Alfazyne l0 mg/kg i.p. (ksilazin)) was used as anesthesia. After that, three layers of polydioxanone sutures were used to close the incision. To lessen postoperative pain, the surgical incisions received 0-5 mL of 0-25 percent bupivacaine and 50 mL/kg of normal saline before and after the procedure. As needed, buprenorphine 0.05 mg/kg was administered for pain.The weight and blood glucose levels of animals that belong to each group were assessed before sacrifice. Glycemia measured with the glucometer (Roche Glucometer Accutrend Plus) expressed in mg/dL. All rats were sacrificed 8 weeks after sleeve gastrectomy operation (fig.1).

**Fig. 1.**
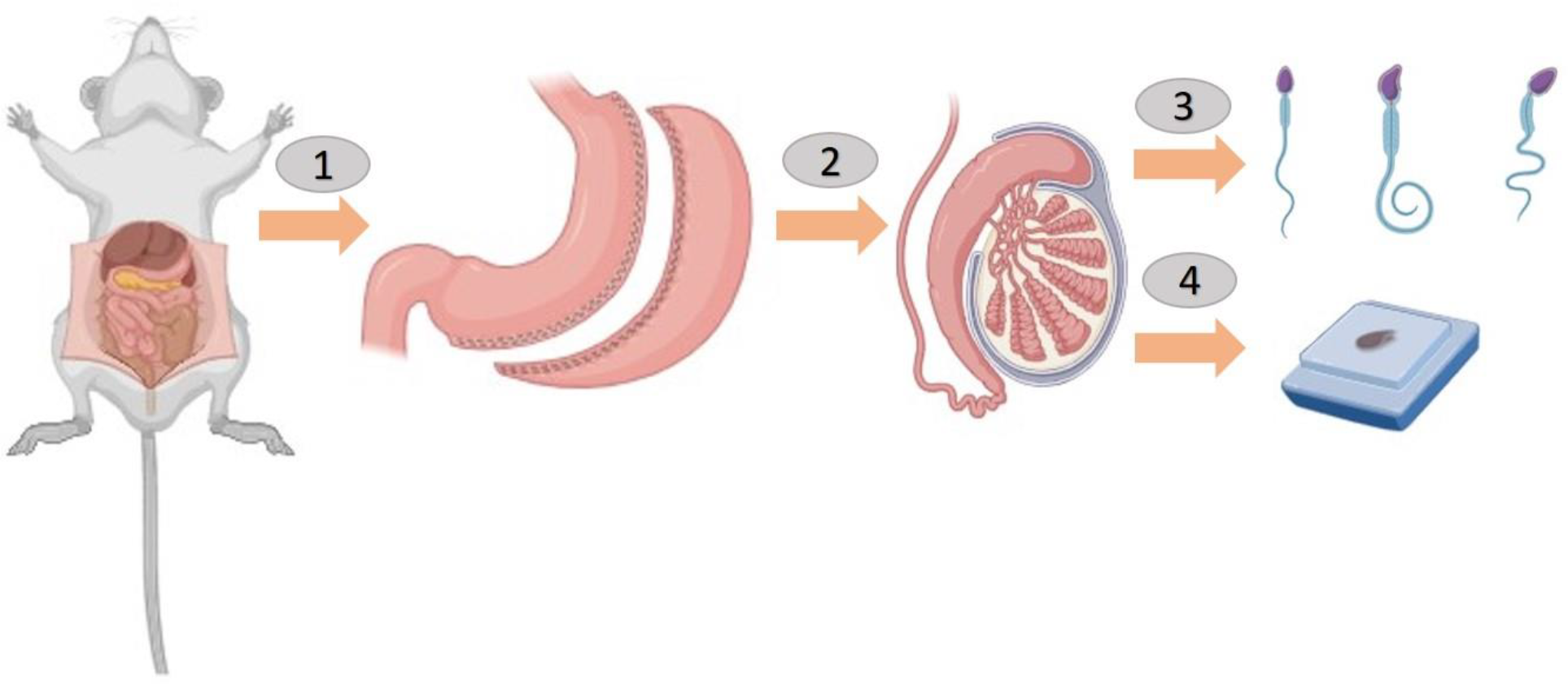
Sleeve gastrectomy surgery was made on rats (1,2) after 8 weeks the rats were anesthetized, their scrotums were opened and the left testicles were removed for sperm count (3) and the right testicles were removed for histology reporting (4).

### Histological examination of testis

Tissue preparation and staining testis specimens were fixed in % 10 tampon formalin and included in paraffin after sacrifice for all groups of rats. All hematoxylin-eosin (HE)-stained slides of testis (20) were analyzed light microscope (Zeiss) both blinded to the group status. Serial sections were cut from each block in the range of 15-30 μm and 5 μm in thickness.

### Evaluation and scoring

#### Assessment of sperm counting and morphological analyze

The left epididymal fluid containing sperm was obtained from all rats as follows. After induction of anesthesia, the epididymis was removed and the tail was separated from the rest of the tissue. Cauda epididymides were minced and placed in a physiological solution adjusted to pH 6.6 and containing 50 mM NaCl, 50 mM K-gluconate, 1.2 mM MgSO4, 0.6 mM CaCl2, 4 mM NaOAc, 1 mM Trisodium citrate, 6.4 mM NaH2PO4, 3.6 mM Na2HPO4 for 15 min at 37°C. Afterward, the remaining tissue was then discarded and the medium was centrifuged at 400 g for 5 min at room temperature to separate the epididymal sperm from the epididymal fluid. Sperm cells were immediately used for the co-incubation assay. After 15 minute spermatozoon falling on all squares in both counting areas counted and observed by light microscopy. The obtained result multiplied by 10^6^ and the number of sperms per milliliter calculated. Spermatozoon morphology (neck and tail) assessment seen with H-E staining [21-23]. Based on these studies, the obtained samples for the evaluation of spermatozoon morphology treated with H-E. On these samples, morphology evaluation performed at light microscope at X400 magnification. 100 spermatozoon in every sample were scored and these evaluated as morphologically normal or abnormal (neck and tail defect) (Fig.1;3). Sperm samples obtained from all groups.

#### Hormone secretion study

At the end of the experiment, the rats dissected under Ketas-Xylazine anesthesia, and blood samples were taken from the intracardiac route were transferred to sterile tubes with anticoagulants. Testosterone kit (Sigma) used to measure testosterone levels and measurements made according to the recommended protocol.

#### Statistical Analysis

Statistical differences between four groups were analysed by either one-way analysis of variance (ANOVA) with Bonferroni’s multiple comparison test (normally distributed data) or Tukey test (non-normally distributed data). Statistical significance was defined as p < 0.05. All analyses were accomplished using the software in GraphPad Prism 3.0 (GraphPad Software, San Diego, CA, USA) and SPSS statistical software version 25 (SPSS Inc., IBM, USA).

## Results

### Diabetes subjects have high glucose levels, which are restored post-sleeve gastrectomy surgery

Eight weeks after sleeve gastrectomy, the diabetes rats weighed significantly less than the DCNT group rats but significantly heavier weight in the CNT groups compared to other all group (fig. 2 A). The blood glucose levels were significantly heavier in DCNT group and given a sleeve gastrectomy (DSG) thus resulted in downregulation of blood glucose levels compared to DSO group (fig. 2B).

**Figure 2.**
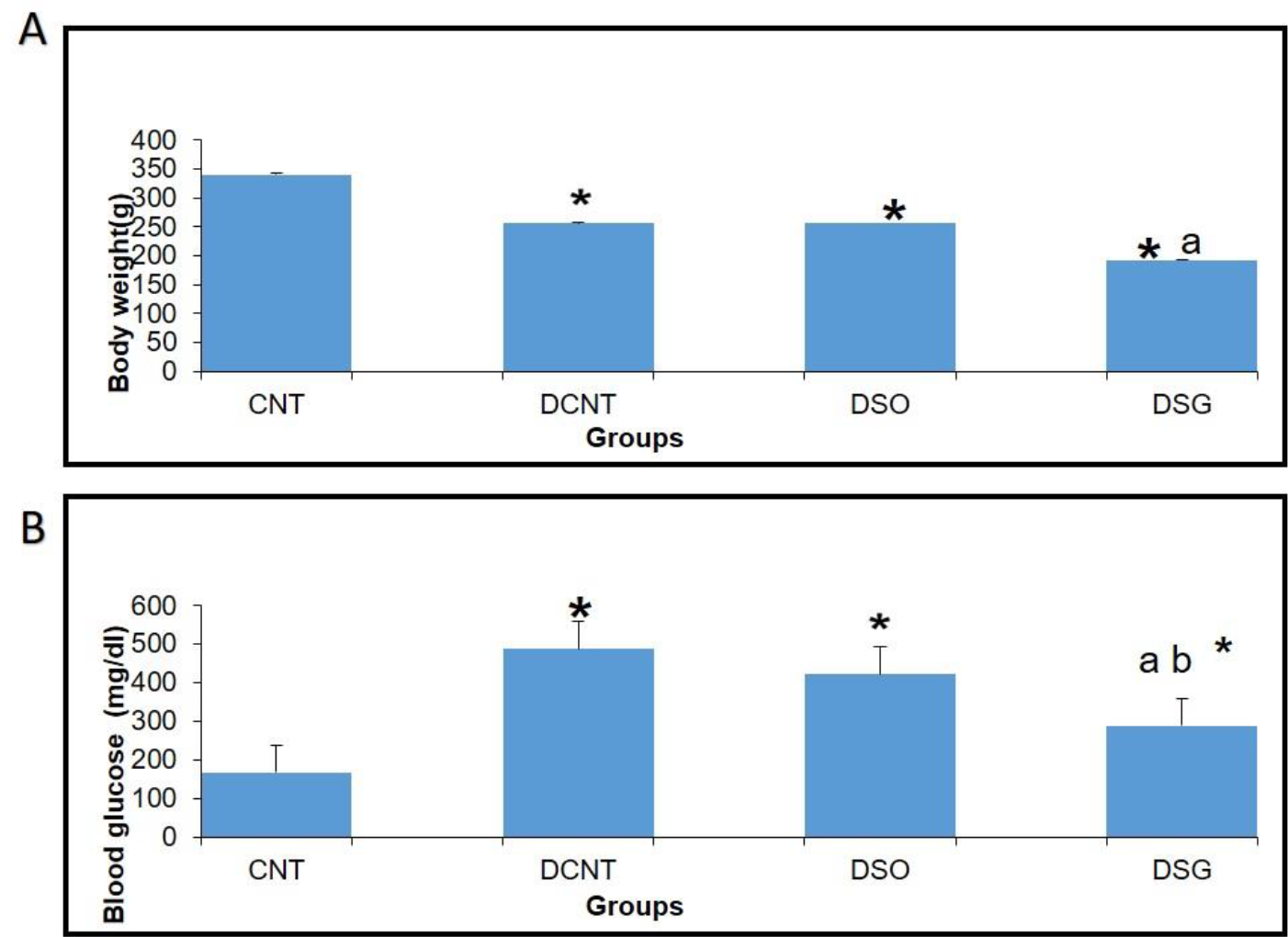
A) Body weights were detected after surgery. * : There is a significant difference between CNT and all other groups (p<0,05). ^a^ : There is a significant difference between DCNT and DSG (p<0,001). ^b^ : There is a significant difference between DSO and DSG groups (p<0,05). B) Concentration of glucose in the blood of four groups after a surgery. * : There is a significant difference between CNT and all other groups (p<0,05). ^a^ : There is a significant difference between DCNT and DSG (p<0,001). ^b^: There is a significant difference between DSO and DSG groups (p<0,001).

### Improvement of rat sperm and testis morphology post-surgery is mediated by testosterone

The testosteron levels were decreased in the diabetes group compared with those in the CNT group (*; p<0,05). On the other hand, after sleeve gastrectomy, the testosterone level was increased significantly compared to the DCNT group (a; p <0.05). After sleeve gastrectomy, testosterone level had increased significantly different from the levels in the DCNT group (a; p<0,05) (Fig.3).

Morphological analysis revealed severely deformity in the seminiferous tubules and disappearance in the seminiferous epithelial cells and enlargement in the interstitial area commonly detected in the DCNT group. In some tubules, it was found that the normal structure of spermatogenic cells preserved (Fig). In the DSO group observed that there was intratubular edema, some tubules did not have germ cells, and loose arrangement of Sertoli and spermatogenic cells (Figure). After sleeve gastrectomy, it was determined that most of the seminiferous tubules, Sertoli cells, and Leydig cells in the interstitial area retain their normal structure. It was evident that the seminiferous tubule lumens were filled with spermium, and the spermium tails were directed towards the lumen. It was noteworthy that all spermatogenic serial cells were present and maintained their integrity (Fig.4A).

Morphology of Spermia; It evaluated according to the neck and tail parts. Hook-shaped normal spermium heads, acrosomes located in the convex region, and centrally located nuclei, neck and tail areas found to be normal. In smear examinations stained with H-E, normal and different morphology spermium observed. In the DCNT group, it determined that the spermatozoa belonging to the DSG group did not have a broken head, and the sperms in the normal morphology similar to the control group were higher (Fig.4B).

Analysis of spermatozoa extracted from the cauda epididymis revealed that the sperm counting was significantly lower in the DCNT and DSO groups than those in the CNT group (*; p<0,001). The sperm counting was higher in the DSG than that in the DSO and DCNT groups (a; p<0,001; b; p<0,05) (Fig.4C). The testes weights were decreased in the DSO and DCNT groups compared with those in the CNT group (*; p<0,05). After sleeve gastrectomy, testes weight had increased compared with those DCNT and DSO groups. So sleeve gastrectomy reversed those changes (a; p<0,05) (Fig 4D).

**Figure 3.**
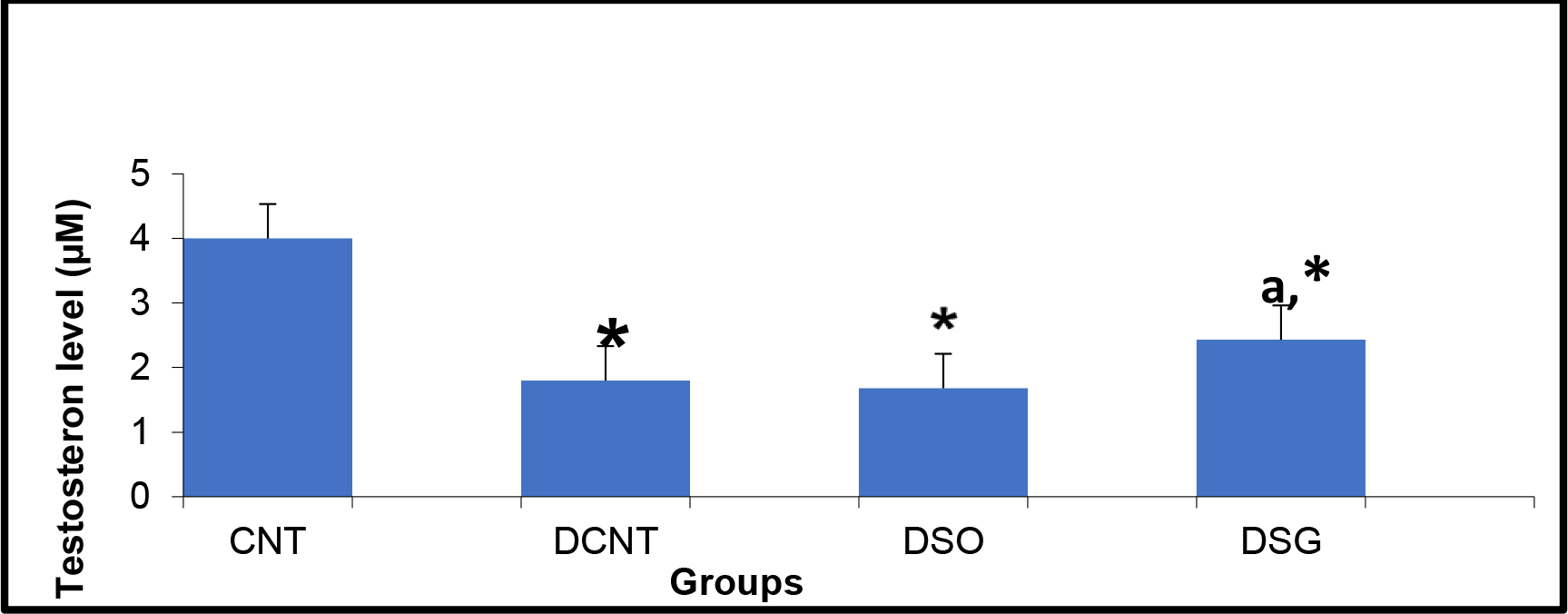
Comparison of testosteron level between the groups at the end of the experiment.

**Figure 4.**
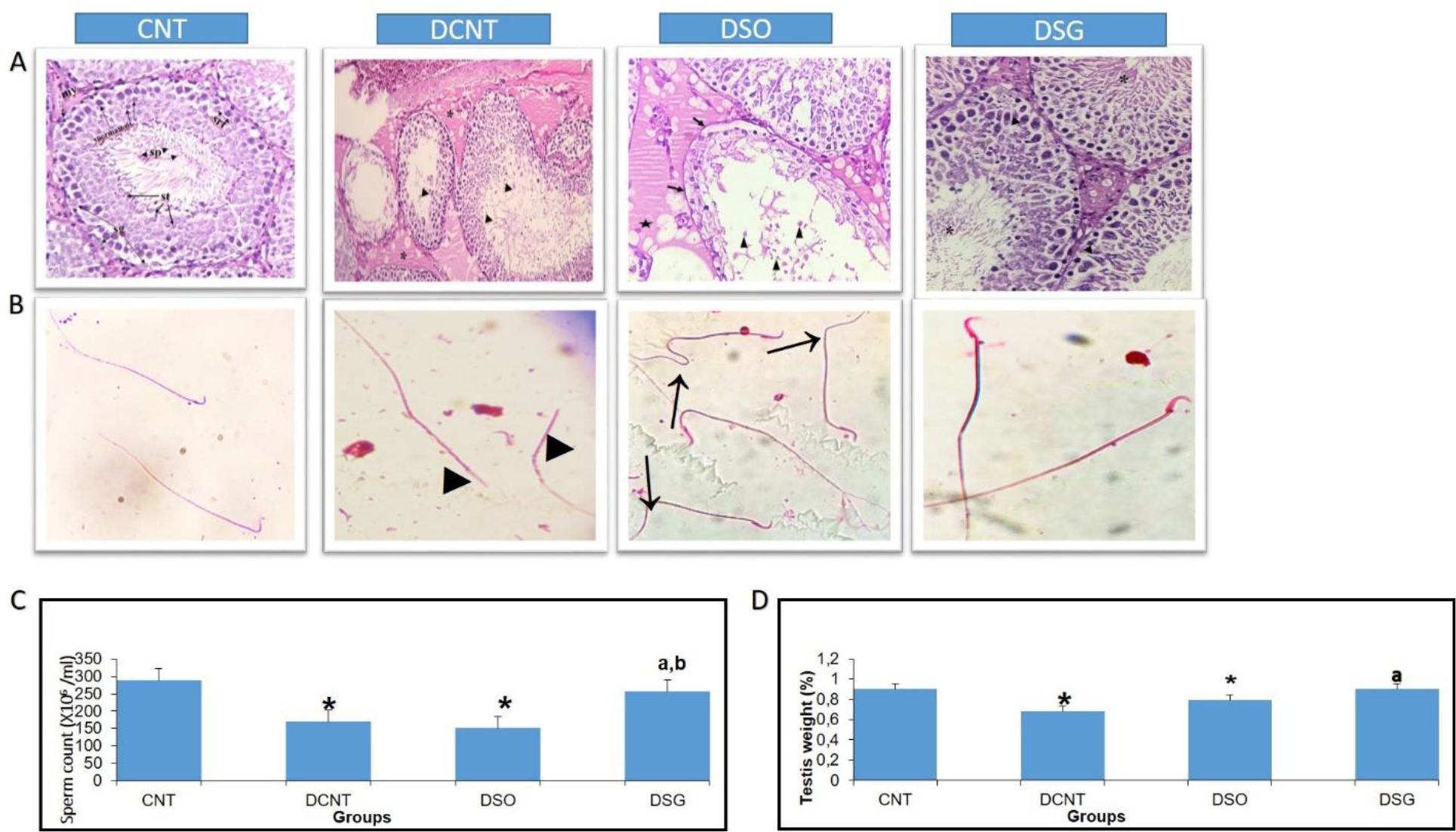
A,B) Representative testis sections and sperm cells staining with H&E bars = 20 μm. C) The findings regarding the sperm counting and D) testicular weights according to the groups.

## Discussion

TIIDM can be seen as a worldwide prevalence of deceleration is between 35-51% [24, 25]. In studies in which experimental TIIDM was created with STZ, it was reported that there was a decrease in the body weight levels of rats compared to the control group [26-32]. In our study, a statistically significant decrease in the body weight of other groups compared to the CNT group was found in accordance with the literature. SG, compared to conventional medical treatment and diet, it has been reported as an effective surgical model for accelerating glucose metabolism and achieving weight loss in both humans [33-36] and rodents [37, 38]. It has been suggested that serum glucose levels and insulin resistance decreased over time in Zucker diabetic obese rat models performed SG compared to controls [37, 38]. In parallel with the studies, in this study sleeve gastrectomy decreased body weight and blood glucose level was statistically compared to the DCNT group(38, 39). In different experimental diabetes model studies, it was found that both body and testicular weight decreased compared to the control group [40-43]. It has been reported that the decrease in testicular weight is caused by deterioration in the structure of the seminiferous tubules, atrophy of the seminiferous tubules and losses in the spermatogenic series [42]. In our study, there was also a decrease in the testicular weight in the diabetes model groups compared to the CNT group. The weight loss in testicular weight results of rats in the DK group was parallel to the studies. In the DSG group, an increase in testicular weight was also observed in parallel with the increase in seminiferous tubule diameter. This increase in testicular weight was correlated with the preservation of the morphology of the cells in the seminiferous tubules. TIIDM affects male fertility in various ways and causes negative effects on the endocrine control of spermatogenesis. As a result of all the biochemical changes observed by diabetes negatively affecting gonadal functions and lowering testosterone levels [41, 42], there is an irregularity in the seminiferous epithelium, an increase in basement membrane thickness and interstitial space. At the same time, vacuolizations in Leydig, germ and Sertoli cells [43], microvascular changes, atrophy of seminiferous tubules, an increase in the number of cells that go to apoptosis in spermatogonium and spermatocytes, as a result of which, a decrease in sperm count has been reported in both experimental animals and humans [44, 45]. A decrease in testicular weight and Leydig cell number are some of the indicators of testicular damage in rat testicles with diabetes model [43, 46]. In our study, it was seen that spermatogenic series cells had a normal appearance in the testicular tissues of rats compared to the CNT group. DCNT and DSO groups subjects showed damage and giant cells in the spermatogenic series, germ cell loss in some tubules, invaginations in the tubule walls, vacuolization in the intertitial area, a decrease in sperm density in the tubule lumen. Giant cells are formed as a result of spermatids merging with each other, and spermatogenic serial cell degeneration is due to the death of Sertoli cells [47]. The study results are consistent with those observations as well as with previous report of a negative effect, as a thickening of tunica albuginea was also detected in the DSO group (46) and a pronounced edema was observed in the interstitial areas between the tubules, which also caused the separation of Leydig cells, and a disorganization of germ cells. Although there have been sperm parameter evaluations in the bariatric surgery studies conducted so far, no studies on testicular tissue morphology have been encountered. The data reported here are not directly comparable with previous data as no study reported the effects of different bariatric surgeries on testicular morphology were revealed for the first time. In our study, dense spermatid was observed in the tubular lumens in the DCNT and DSO groups compared to the DSG group. It was found that the spermatogenic cell series in the seminiferous tubule wall also maintained their normal structure. No pathological condition was detected in the Leydig cells located in the interstitial space. Sleeve gastrectomy reversed the spermatogenic series and edema in testicular tissues.It is known that differences in seminiferous tubule diameters occur as a result of histopathological changes in testicular tissues and losses in spermatogenic cell series in STZ-induced diabetes models [42-48]. In this study results are consistent with previous reports a decrease in the seminiferous tubule diameter was found in the DCNT and DSO groups compared to the CNT group. After sleeve gasrectomy was found to have an increasing effect on this decrease in seminiferous tubule diameters also high sperm count, which increased along with the density of germ cells and spermatids present in the lumen, was interpreted as the reasons for the increase in tubule diameter. When we look at the seminiferous tubule diameters, it is thought that the germ cell loss observed in the DCNT and DSO groups and the absence of spermatid cells in the lumen due to this loss reduce the seminiferous tubule diameters.

Previous studies showed that the effects of diabetes on sperm parameters have reported changes such as decreased sperm count and motility [32, 41], pathological sperm morphology [41, 49] and increased semen volume [46] or decreased [25], penile erection and ejaculation deterioration [24]. Another study conducted among men with diabetes found an increase in sperm concentration and total sperm production, deceleration in motility, while it did not find any differences in sperm morphology [50]. Data obtained from experimental study models; It is suggested that TIIDM damages many male reproductive functions such as sperm concentration, motility and morphology [51, 52]. Due to the increasing incidence of morbid obesity, bariatric surgery, which has a long-term effectiveness on weight loss, comorbidities and mortality, has become an increasingly common form of treatment, especially in China [53-56]. Therefore, it also closely concerns young patients of reproductive age. Therefore, the possible effects of bariatric surgeries on semen quality and male fertility are of clinical importance. However, the results in the literature show conflicting results, especially regarding the effects of bariatric surgical procedures on semen parameters [7, 57, 58]. Some researchers suggest that improvement in sexual life quality [58] and hormonal profile improvements can be expected after bariatric operations [57, 58]. While some studies did not find a significant difference in sperm parameters after bariatric surgery [7], Di Frega et al. observed secondary azoospermia in patients after surgery [56]. Sermondade et al. [57] reported that 3 months after surgery, there was a serious deterioration in semen parameters in three patients, but it reversed after 24 months (including oligoasthenoteratozoospermia). El Bardisi et al. reported that sex hormone profile and semen characteristics of 46 patients before and after SG. It has been reported that there is a tendency towards testosterone increase and semen recovery in the short term, but no results regarding long-term effects have been obtained [10]. According to Lazaros et al., reported that sperm parameters in two male patients before and after bariatric surgery, and a significant decrease in parameters was recorded 12 to 18 months after surgery. With this result, they came to the conclusion that bariatric surgery can negatively affect spermatogenesis in the first months after surgery [59]. To the best of our knowledge, the data reported here directly comparable with previous data a decrease in seminiferous tubule diameter, a low sperm count and abnormal sperm morphology were detected along with seminiferous tubule damage in the DCNT and DSO groups compared to the CNT group. After sleeve gastrectomy spermatozoon morphology similar to compared with those in the CNT group Our results were indicating that sleeve gastrectomy may have positive effects on male reproductive potential in the first months after surgery. It has been shown that there is a spermatozoon pathology in the diabetes model, and this pathology is also seen in the tail section [54]. Analysis of spermatozoa extracted from the cauda epididymis revealed that tail pathology was found in the DCNT and DSO groups. Sleeve gastrectomy reversed those changes. Sleeve gastrectomy thus improved sperm quality and increased reproductive capacity. Endocrine dysregulation is another characteristic typical of obesity, with declines in serum levels of total and free testosterone, LH, FSH, and increase in estradiol (E2) levels accompanying the accumulation of fatty tissue [60-62]. The study results are consistent with those observations as well as with previous reports of a negative effect of estrogen on the hypothalamus through pulsed regulation of gonadotropin-releasing hormone (GnRH). In those studies, increases in estrogen elicited decreases of GnRH, suppression of both LH and FSH secretion, and reduced testosterone secretion and spermatozoa production [63, 64]. In this rat model, sleeve gastrectomy decreased E2 expression in addition to resulting in weight loss. There are also clinical studies conducted in parallel with experimental studies [7] that show a decrease in testosterone, LH, FSH levels in diabetes [65]. A decrease in testosterone levels is one of the causes of testicular damage due to diabetes. Many researchers suggest that the decrease in serum testosterone levels in TIIDM is caused by steroidogenetic damage to Leydig cells [57]. In our study, the testosterone levels of the diabetes model rats decreased statistically compared to the CNT group. It is suggested that there are increases in total and free testosterone, SHBG, LH and FSH in men who have undergone bariatric surgery. In parallel with the improvements in male sex hormone levels, a significant increase in erectile function after bariatric surgery is also known [66, 67]. The improvement in total testosterone plasma levels 1 month after bariatric surgery is reported in different studies [68-70]. The study results are consistent with those observations as well as with previous reports of positive effect of Sleeve gastrectomy increased testosterone level [66, 67].

When our findings were evaluated, it was interpreted that SG surgery decelerated the negative effects of diabetes on male fertility in the short term among the surgical techniques we applied. As a result, it was found that the diabetes model negatively affects the spermatogenic series and sperm morphology by affecting the testicles, but sleeve gastrectomy, a type of bariatric surgery, reduces blood glucose levels and corrects testicular tissue integrity and sperm morphology. In individuals with TIIDM, it is recommended to plan/ conduct prospective studies to assess whether bariatric surgery models are effective on sperm parameters and male fertility and their long-term effects.

## Author contributions

Gamze Tumentemur had full access to all the data in the study and takes responsibility for the integrity of the data and the accuracy of the data analysis.

*Study concept and design:* Gamze Tumentemur

*Acquisition of data:* Gamze Tumentemur, Mustafa Titiz

*Analysis and interpretation of data:* Gamze Tumentemur, Mustafa Titiz

*Drafting of the manuscript:* Gamze Tumentemur,

*Statistical analysis:* Gamze Tumentemur,

*Obtaining funding:* None.

*Administrative, technical, or material support:* None.

*Supervision:* Alev Bobus

## REFERENCES

[1] Berghofer, A., Pischon, T., Reinhold, T., Apovian, C.M., Sharma, A.M., Willich, S.N. (2008). Obesity prevalence from a European perspective: a systematic review. BMC Public Health, 8, 200–215.

[2] La Vignera, S., Calogero, A.E. Condorelli, R. ve diğerleri. (2009). Andrological characterization of the patient with diabetes mellitus. Minerva Endocrinol, 34, 1–9.

[3] Jangir, R.N., Jain, G.C. (2014). Diabetes mellitus induced impairment of male reproductive functions: a review. Curr. Diabetes Rev, 10, 147–157.

[4] Dias, T. R., Alves, M. G., Silva, B. M., & Oliveira, P. F. (2014). Sperm glucose transport and metabolism in diabetic individuals. Molecular and cellular endocrinology, 1, 37–45.

[5] Ballester, J., Muñoz, M. C., Domínguez, J., Rigau, T., Guinovart, J. J., & Rodríguez-Gil, J. E. (2004). Insulin-dependent diabetes affects testicular function by FSH and LH-linked mechanisms. Journal of andrology, 5, 706–719.

[6] Dall, T., Edge, M. S., Zhang, Y., Martin, J., Chen, Y., Hogan, P. (2007). Economic costs of diabetes in the U.S. Diabetes Care, 31, 596–615

[7] Reis, L.O., Zani, E.L., Saad, R.D. ve diğerleri. (2012). Bariatric surgery does not interfere with sperm quality–a preliminary long-term study. Reprod Sci, 19, 1057–6.

[8] Lazaros, L., Hatzi, E., Markoula, S., Takenaka, A. ve diğerleri. (2012). Dramatic reduction in sperm parameters following bariatric surgery: report of two cases. Andrologia, 6, 428–32.

[9] Legro, R.S., Kunselman, A.R., Meadows, J.W., Kesner, J.S., Krieg, E.F., Rogers, A.M. (2015). Time related increase in urinary testosterone levels and stable semen analysis parameters after bariatric surgery in men. Reprod Biomed Online, 2, 150–6.

[10] El Bardisi, H., Majzoub, A., Arafa, M., AlMalki, A., Al Said, S., Khalafalla, K. ve diğerleri. (2016). Effect of bariatric surgery on semen parameters and sex hormone concentrations: a prospective study. Reprod Biomed Online, 5, 606–11.

[11] Porries, W. J., Swanson, M. S., Macdonald, K. G., Long, S. B., Morris, P. G. ve diğerleri. (1995). Who would have thought it? An operation proves to be the most effective therapy for adult-onset diabetes mellitus. Annals of Surgery, 222, 339–352.

[12] Sjoström, L., Narbro, K., Sjostrom, C.D., Karason, K., Larsson, B., Wedel, H., Lystig, T., Sullivan, M., Bouchard, C., Carlsson, B., Bengtsson, C., Dahlgren, S., Gummesson, A., Jacobson, P., Karlsson, J., Lindroos, A.K., Lonroth, H., Naslund, I., Olbers, T., Stenlof, K., Torgerson, J., Agren, G., Carlson, L.M. (2007). Effects of bariatric surgery on mortality in Swedish obese subjects. N. Engl J Med, 357, 741–752.

[13] Buchwald, H., Estok, R., Fahrbach, K., Banel, D., Jensen, M.D., Pories, W.J. ve diğerleri. (2009). Weight and type 2 diabetes after bariatric surgery: systematic review and meta analysis. Am J Med, 122, 248–256.

[14] Glatt, D., Sorenson, T. (2011). Metabolic and bariatric surgery for obesity: a review. SD Med Spec 57–62.

[15] Brown, W., Burton, P., Anderson, M., Korin, A., Dixon, J., Hebbard, G. ve diğerleri. (2008). Symmetrical pouch dilatation after laparoscopic adjustable gastric banding: incidence and management. Obes Surg, 18, 1104–1108.

[16] Hutter, M.M., Schirmer, B.D., Jones, D.B., Ko, C.Y., Cohen, M.E., Merkow, R.P. ve diğerleri. (2011). First report from the American College of Surgeons Bariatric Surgery Center Network: laparoscopic sleeve gastrectomy has morbidity and effectiveness positioned between the band and the bypass. Ann Surg, 254, 410–420.

[17] Pech, N., Meyer, F., Lippert, H., Manger, T., Stroh, C. (2012). Complications and nutrient deficiencies two years after sleeve gastrectomy. BMC Surg, 12, 13.

[18] Karaca, T., Demirtaş, S., Karaboğa, İ., Ayvaz, S. (2015). Protective effects of royal jelly against testicular damage in streptozotocin-induced diabetic rats. 1, 27 –32.

[19] Masiello, P., Broca, C., Gross, R., Roye, M., Manteghetti, M., Hillaire-Buys, D., Novelli, M., Ribes, G. (1998). Experimental NIDDM: development of a new model in adult rats administered streptozotocin and nicotinamide. Diabetes, 2, 224–9.

[20] Sánchez, R.; Villagrán, E.; Risopatrón, J. & Célis, R. (1994). Evaluation of nuclear maturity in human spermatozoa obtained by sperm preparation methods. Andrologia, 3, 173–6.

[21] Aksoy, E., Aktan, T. M., Duman, S., Cuce, G. (2012). Assessment of Spermatozoa Morphology under Light Microscopy with Different Histologic Stains and Comparison of Morphometric Measurements. Int J Morphol, 4, 1544–1550.

[22] Wild, S., Roglic, G., Green, A., Sicree, R., King, H. (2004). Global prevalence of diabetes: estimates for the year 2000 and projections for 2030. Diabetes Care, 5, 1047–1053.

[23] La Vignera, S., Calogero, A.E., Condorelli, R., Lanzafame, F., Giammusso, B., Vicari, E. (2009). Andrological characterization of the patient with diabetes mellitus. Minerva Endocrinol, 1, 1–9.

[24] Singh, K., Devi, S., Pankaj, P. P. (2016). Diabetes Associated Male Reproductive Dysfunctions: Prevalence, Diagnosis and Risk Factors. Int J Drug Dev & Res, 8, 007–010.

[25] Bener, A., Al-Ansari, A.A., Zirie, M., Al-Hamaq, A.O. (2009). Is male fertility associated with type 2 diabetes mellitus. Int Urol Nephrol, 4:777–84.

[26] Ünlüçerçi, Y., Bekpınar, S., Koçak, H. (2000). Testis Glutathione Peroxidase and Phospholipid Hydroperoxide Glutathione Peroxidase Activities in AminoguanidineTreated Diabetic Rats. Academic Press, 379, 217–220.

[27] Andallu, B., Varadacharyulu, N.C. (2003). Antioxidant role of mulberry [Morus indica L. cv. Anantha] leaves in streptozotocin-diabetic rats. Clinica Chimica Acta, 338, 3–10.

[28] Aksoy, N., Vural, H., Sabuncu, T., Aksoy, S. (2003). Effects of melatonin on oxidative antioxidative status of tissues in streptozotocin-induced diabetic rats. Cell Biochem Funct, 21, 121–125.

[29] Ricci, G., Catizone, A., Esposito, R., Pisanti, F.A., Vietri, M.T., Galdieri, M. (2009). Diabetic rat testes: morphological and functional alterations. Andrologia, 41, 361–368.

[30] Guneli, E., Tugyan, K., Ozturk, H., Gumustekin, M., Cilaker, S., Uysal, N. (2008). Effect of Melatonin on Testicular Damage in Streptozotocin-Induced Diabetes Rats. Eur Surg Res, 40, 354–360.

[31] Buchwald, H., Estok, R., Fahrbach, K. ve diğerleri. (2009). Weight and type 2 diabetes after bariatric surgery: systematic review andmeta-analysis. Am J Med, 3, 248–56.

[32] Khaki, A., Nouri, M., Fathiazad, F., Ahmadi-Ashtiani, H., Rastgar, H., & Rezazadeh, S. (2009). Protective effects of quercetin on spermatogenesis in streptozotocin-induced diabetic rat. Journal of Medicinal Plants, 29, 57–64.

[33] Eid, G.M., Brethauer, S., Mattar, S.G., Titchner, R.L., Gourash, W. ve Schauer, P.R. (2012). Laparoscopic sleeve gastrectomy for super obese patients: forty-eight percent excess weight loss after 6 to 8 years with 93% follow-up. Ann Surg, 256, 262–265.

[34] Rosenthal, R.J., Diaz, A.A., Arvidsson, D., Baker, R.S., Basso, N., Bellanger, D., Boza, C., El Mourad, H. ve diğerleri. (2012). International sleeve gastrectomy expert panel consensus statement: best practice guidelines based on experience of >12,000 cases. Surg Obes Relat Dis, 8, 8–19.

[35] Barzin, M., Motamedi, M.A., Serahati, S., Khalaj, A., Arian, P. ve diğerleri. (2017). Comparison of the effect of gastric bypass and sleeve gastrectomy on metabolic syndrome and its components in a cohort: Tehran obesity treatment study [TOTS]. Obes Surg, 27, 1697–1704.

[36] Mason, M.C. ve Pournaras, D.J. (2017). Sleeve gastrectomy: Managing the morbidity of obesity. Angiology, 3319717724941.

[37] Kadera, B., Portenier, D., Yurcisin, B., Demaria, E., Gaddor, M. and Jain-Spangler, K. (2013). Evidence for a metabolic mechanism in the improvement of type 2 diabetes after sleeve gastrectomy in a rodent model. Surg Obes Relat Dis, 9, 447–452.

[38] Ying, F., Chen, Z., Jianxiang, N., Lin, Z., Yang, Z., Wen, W., Zelan, H., Hui, W., Ping, H., Qinjie, N., JingJing, X., Jing, Z. (2018). Effects of sleeve gastrectomy on lipid and energy metabolism in ZDF rats via PI3K/AKT pathway Am J Transl Res, 11, 3713–3722.

[39] Dong, S., Shaozhuang, L., Guangyong, Z., Punsiri, C., Chunxiao, H., Haifeng, H., Mingxia, L., Sanyuan, H. (2014). Sub-sleeve gastrectomy achieves good diabetes control without weight loss in a non-obese diabetic rat model. Surg Endosc, 28, 1010–1018.

[40] Prabhakara, P.K., Prasad, R., Ali, S., Doblec, M. (2013). Synergistic interaction of ferulic acid with commercial hypoglycemic drugs in streptozotocin induced diabetic rats. Phytomedicine, 6, 488–494.

[41] Steger, R.W., Rabe, M.B. (1997). The effect of diabetes mellitus on endocrine and reproductive function. Proc Soc Exp Biol Med, 1: 1–11.

[42] Özcan, Ö. (2017). Streptozotosin (stz) ile diyabet oluşturulmuş sıçanlarda kuersetin“in testis dokusuna etkisi. Yüksek Lisans Tezi, Eskişehir Osman Gazi Üniversitesi Sağlık Bilimleri Enstitüsü.

[43] Kianifard, D., Sadrkhanlou, R. A., & Hasanzadeh, S. (2012). The ultrastructural changes of the sertoli and leydig cells following streptozotocin induced diabetes. Iranian journal of basic medical sciences, 1, 623–635.

[44] Türkiye Diyabet Vakfı Yayınları (2013). Diyabet Tanı ve Tedavi Rehberi. 3. Baskı, Armoni Nüans Baskı Sanatları, İstanbul.

[45] Türkiye diyabet tanı ve tedavi rehberi, (2019). İstanbul.

[46] Cameron DF, Murray FT, Drylie DD. (1985). Interstitial compartment pathology and spermatogenic disruption in testes from impotent diabetic men. The Anatomical Record, 1, 53–62.

[47] Kaya, M. (1986). Sertoli cells and various types of multinucleates in the rat seminiferous tubules following temporary ligation of the testicular artery. Journal of Anatomy, 144, 145.

[48] Cai, L., Chen, S., Evans, T., Deng, D. X., Mukherjee, K., & Chakrabarti, S. (2000). Apoptotic germ-cell death and testicular damage in experimental diabetes: prevention by endothelin antagonism. Urological research, 5, 342–347.

[49] Jangir, R.N., Jain, G.C. (2014). Diabetes mellitus induced impairment of male reproductive functions: a review. Curr. Diabetes Rev, 10, 147–157.

[50] Oufi, H.G., Al-Shawi, N.N. (2014). The effects of different doses of silibinin in combination with methotrexate on testicular tissue of mice. Eur J Pharmacol, 730, 36–40.

[51] Aksoy, E., Aktan, T. M., Duman, S., Cuce, G. (2012). Assessment of Spermatozoa Morphology under Light Microscopy with Different Histologic Stains and Comparison of Morphometric Measurements. Int J Morphol, 4, 1544–1550.

[52] Scarano, W.R., Messias, A.G., Oliva, S.U., Klinefelter, G.R., Kempinas, W.G. (2005). Sexual behaviour, sperm quantity and quality after short-term streptozotocin-induced hyperglycaemia in rats. Int J Androl, 4, 482–8.

[53] Handelsman, D.J., Conway, A.J., Boylan, L.M., Yue, D.K. and Turtle, J.R. (1985). Testicular function and glycemic control in diabetic men. A controlled study. Andrologia, 17, 488–496.

[54] Mingrone, G., Panunzi, S., De Gaetano, A., Guidone, C., Laconelli, A. ve diğerleri. (2012). Bariatric Surgery versus Conventional Medical Therapy for Type 2 Diabetes. N Engl J Med, 17, 1577–1585.

[55] Du, X., Dai, R., Zhou, H.X. ve diğerleri. (2016). Bariatric surgery in China: How is this new concept going? Obes Surg, 26, 2906–12.

[56] Di Frega, A.S., Dale, B., Di Matteo, L., Wilding, M. (2005). Secondary male factor infertility after Roux-enY gastric bypass for morbid obesity: case report. Hum Reprod, 4 997.

[57] Sermondade, N., Massin, N., Boitrelle, F., Pfeffer, J., Eustache, F., Sifer, C. ve diğerleri. (2012). Sperm parameters and male fertility after bariatric surgery: three case series. Reprod BiomedOnline, 24, 206–210.

[58] Legro, R.S., Kunselman, A.R., Meadows, J.W., Kesner, J.S., Krieg, E.F., Rogers, A.M. (2015). Time related increase in urinary testosterone levels and stable semen analysis parameters after bariatric surgery in men. Reprod Biomed Online, 2, 150–6.

[59] Lazaros, L., Hatzi, E., Markoula, S., Takenaka, A. ve diğerleri. (2012). Dramatic reduction in sperm parameters following bariatric surgery: report of two cases. Andrologia, 6, 428–32.

[60] King, P., Peacock, I., Donnelly, R. (1999). The UK Prospective Diabetes Study (UKPDS): clinical and therapeutic implications for type 2 diabetes. British Journal of Clinical Pharmacology, 5, 643–8.

[61] Peynirci, H. (2010). Deneysel diabetik nefropatide irbesartan, antioksidan ve kombinasyon tedavilerinin karşılaştırılması. Trakya Üniversitesi Tıp Fakültesi İç Hastalıkları Anabilim Dalı. Uzmanlık Tezi, Edirne.

[62] Abacı, A., Böber, E., Büyükgebiz, A. (2007). Tip 1 diabet. Güncel Pediatri, 5, 1–10.

[63] Türkiye diyabet tanı ve tedavi rehberi, (2019). İstanbul.

[64] Zimmet, P., Tuomi, T., Mackay, R., Rowlwy, M., Knowles, W., Cohen, M., ve diğerleri.

[65] Schauer, P.R., Mingrone, G., Ikramuddin, S., Wolfe, B. (2016). Clinical Outcomes of Metabolic Surgery: Efficacy of Glycemic Control, Weight Loss, and Remission of Diabetes. Diabetes Care, 39, 902–11.

[66] Botella-Carretero, J.I., Balsa, J.A., Gómez-Martin, J.M., Peromingo, R., Huerta, L., Carrasco, M., Arrieta, F., Zamarron, I. ve diğerleri. (2013). Circulating free testosterone in obese men after bariatric surgery increases in parallel with insulin sensitivity. J Endocrinol Invest, 4, 227–32.

[67] Dhindsa, S., Prabhakar, S., Sethi, M., Bandyopadhyay, A., Chaudhuri, A., Dandona, P. (2004). Frequent occurrence of hypogonadotropic hypogonadism in type 2 diabetes. J Clin Endocrinol Metab, 11, 5462–8.

[68] Weiss, H.G., Nehoda, H., Labeck, B., Hourmont, K., Marth, C., Aigner, F. (2001). Pregnancies after adjustable gastric banding. Obes Surg, 11, 303–306.

[69] Yong, W., Quanbing, C., Wenhui, Q. (2018). Effect of Bariatric Surgery on Semen Parameters: A Systematic Review and Meta-Analysis. Med Sci Monit Basic Res, 24, 188–197.

[70] Di Vincenzo, A., Busetto, L., Vettor, R., Rossato, M. (2018). Obesity, Male Reproductive Function and Bariatric Surgery. Front Endocrinol, 9, 769.

